# One-Pot NADH-Mediated Physiological Redox-Controlled Synthesis of Papain-Stabilized Copper Nanoclusters with Preserved Bioactivity for Efficient Drug Delivery

**DOI:** 10.64898/2026.04.20.719779

**Authors:** Aman Singh, Ananya Anand, Vanshika, Anujit Balo, Farhan Ahmed, Mirzada Mohd Qounane, Koyel Banerjee-Ghosh, Ritaban Halder, Shirsendu Ghosh

**Affiliations:** Departments of Chemistry, Indian Institute of Technology, Jammu, NH-44, PO Nagrota, Jagti, Jammu and Kashmir 181221 India; Departments of Bioscience & Bioengineering, Indian Institute of Technology, Jammu, NH-44, PO Nagrota, Jagti, Jammu and Kashmir 181221 India; Department of Chemistry, Indian Institute of Technology Hyderabad, Telangana 502284, India; Department of Life Sciences, Gandhi Institute of Technology and Management, Hyderabad Campus, Rudraram, Telangana 502329, India; Department of Chemistry, University of Southern California, Los Angeles, California 90089-1062, United States; Past Address: Department of Chemistry, Gandhi Institute of Technology and Management, Hyderabad Campus, Rudraram, Telangana 502329, India

**Keywords:** Protein-Stabilized Copper Nanoclusters, NADH-Mediated Reduction, Biocompatible Synthesis, Protein Conformational Stability, Drug Delivery Nanoplatform, Molecular Dynamics Simulation

## Abstract

Protein-protected copper nanoclusters are promising candidates for bioimaging and therapeutic applications. However, harsh reducing conditions required for their synthesis, compromising the protein’s structural integrity and bioactivity. Here, we report a one-pot, physiologically compatible aqueous synthesis for papain-protected copper nanoclusters (Pap-CuNCs) with blue photoluminescence and excellent photostability using nicotinamide adenine dinucleotide (NADH) as a biological reducing agent. The mild conditions (ambient temperature, neutral pH) enable the simultaneous formation of a metallic Cu^0^ core while stabilizing the helical content of the protein. This approach introduces a physiologically redox-controlled strategy for nanocluster formation, establishing physiological redox chemistry as a governing principle for controlling nanoscale structure and protein conformational stability.

Spectroscopic and microscopic studies have demonstrated the presence of crystalline nanoclusters with a protein corona that undergoes α-helical stabilization as revealed by circular dichroism. Notably, atomistic simulation studies reveal preferential binding of the copper core in the protein’s active site, enhancing the α-helical content of papain, consistent with experimental observations.

Functionally, the Pap–CuNCs possess biocompatibility and serve as an effective delivery platform for 5-fluorouracil, leading to a 50-fold decrease in IC50 for HeLa cells without causing cytotoxicity to normal cells. This establishes a generalizable framework for bio-integrated nanocluster design under biologically compatible conditions.

**TOC Text:** 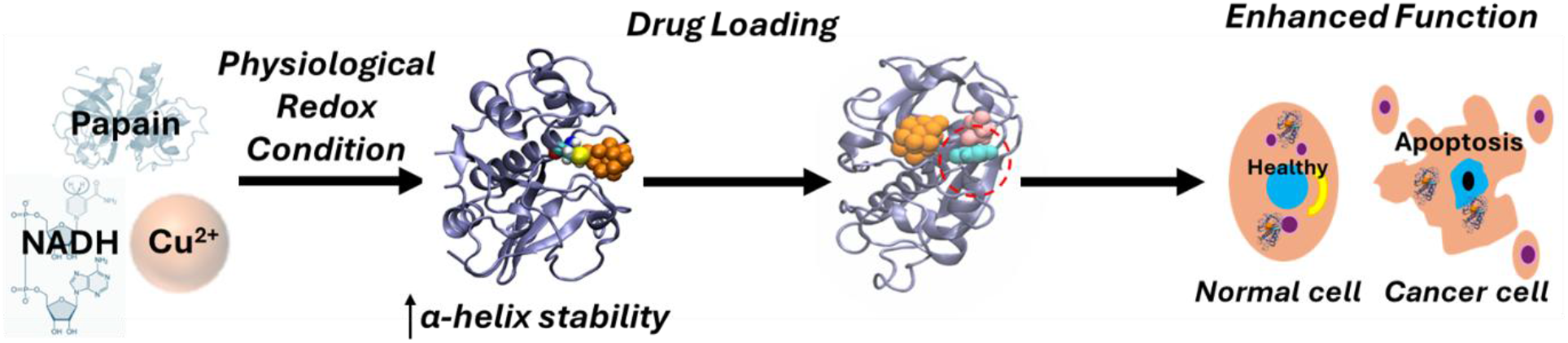

NADH-mediated physiological redox synthesis of papain-stabilized copper nanoclusters with enhanced α-helical stability. This bio-integrated nanoplatform facilitates drug loading and delivery, resulting in a ∼50-fold increase in cytotoxicity in cancer cells while maintaining excellent biocompatibility toward normal cells.

## 1. Introduction

Ligand-protected metal nanoclusters (CuNCs) consist of an ultrasmall inorganic core stabilized by an organic shell (colloquially dubbed as ‘corona’).^[1,2]^ Their core dimension (<3 nm)^[1–3]^ is comparable to the Fermi wavelength of electrons^[2,4]^, which induces molecule-like electronic transitions and discrete energy levels, distinguishing them from their bulk metallic counterparts^[1–5]^. Among coinage metals, gold^[6]^ and silver^[5,7]^ nanoclusters have been thoroughly investigated due to their high stability in the metallic state and well-defined photophysical behavior. In the recent decade, CuNCs have emerged as one of the key members due to their chemical similarity with silver nanoclusters (AgNCs) and gold nanoclusters (AuNCs), different fluorescence properties and most importantly the availability of low-cost and earth-abundant precursors^[8–14]^.

Copper also plays a crucial biological role as a cofactor for the biological oxidation/reduction reaction enzymes, such as cytochrome c oxidase and lysyl oxidase.^[15,16]^ Due to easier metabolism and efficient chelating process in cases of excess accumulation, copper is considered more bio-compatible compared to gold and silver.^[11,15,16]^ Thus, CuNCs are regarded as promising candidates for bioimaging, biosensing, and drug delivery applications, potentially serving as alternatives to conventional quantum dots. However, a significant challenge remains in stabilizing CuNCs, as copper is more susceptible to oxidation and redox-mediated degradation than other noble metal counterparts, often requiring aggressive reductive conditions that are less biocompatible.^[9]^

A broad range of organic ligands, including dendrimers, poly(acrylic acid) polymer, as well as biomolecules such as peptides, DNA, RNA, and proteins, have been employed as stabilizing and/or reducing agents to synthesize CuNCs.^[9]^ Proteins attract particular attention because the protein corona not only stabilizes the metallic core through coordination of functional groups (–NH2, –COOH, –OH, and –SH) but also imparts intrinsic biological functionality. Although a substantial number of reports exist on protein-stabilized CuNCs, the synthesis protocols include either strong inorganic reducing agents (e.g., hydrogen peroxide^[17,18]^ or sodium borohydride^[19]^ or hydrazine hydrate^[11,20,21]^), highly alkaline conditions^[8]^, or elevated temperatures^[22]^, leading to protein unfolding, diminishing bioactivity, and limiting translational potential. Thus, there is an urgent need for a physiologically compatible synthetic method that enables the formation of nanoclusters with the protein structure and function intact.

Papain, a cysteine protease with commercial availability at a low cost, offers a promising, though less explored, protein template for CuNC stabilization.^[21,23,24]^ Besides its proteolytic activity, papain has shown selective apoptosis-inducing activity toward cancer cells, with minimal cytotoxic effects on normal cells.^[25,26]^ Additionally, papain has shown potential in wound debridement, the enhancement of the rate of transdermal drug absorption, and the reduction of mucus viscosity.^[27]^ The various biological activities of papain have shown the potential for the use of the enzyme, not only as a template in the preparation and stabilization of CuNCs but also as a bioactive interface in nanomedicine^[28]^. However, the use of papain in CuNCs is still in its infancy, with only one report being limited to the use of the enzyme in the sensing of hydrogen peroxide and the antimicrobial activity of the CuNCs^[21]^. Therefore, a method for the preservation of the bioactivity of papain in the preparation and stabilization of CuNCs is a potential route to the diverse therapeutic uses of the enzyme.

In biological systems, redox chemistry is tightly controlled in mild, near-neutral conditions. Nicotinamide adenine dinucleotide (NADH) is a ubiquitous biological reductant in glycolysis, the Krebs cycle, and electron transport.^[29–31]^ NADH is a fascinating reductant because it is more stable at physiological pH (∼7) and at room temperature (∼25 °C) compared to sodium borohydride, whose half-life at these conditions is on the order of seconds. The use of NADH as a biocompatible reductant, therefore, provides a promising route for synthesizing protein-protected metal nanoclusters under conditions compatible with protein structural integrity.

Despite significant progress, a general strategy that couples nanocluster formation with preservation of protein structure under physiological redox conditions remains elusive. In this context, we propose that physiological redox chemistry can act as a governing factor in simultaneously directing nanocluster formation and modulating protein conformational stability. Herein, we report a one-pot, fully aqueous, physiologically compatible synthesis of blue-emitting copper nanoclusters stabilized by papain (Pap–CuNCs) using NADH as a biological reducing agent. According to the best of our knowledge, this represents one of the first demonstrations of use of NADH, for the synthesis of protein-coated metal nanoclusters. The synthesis proceeds at ambient temperature and near-neutral pH, enabling preservation of protein functionality. The resulting Pap–CuNCs exhibit strong photoluminescence with a notable quantum yield (∼18%) and high photostability. Comprehensive spectroscopic and microscopic analyses confirm the formation of metallic face-centered cubic (FCC) Cu^0^ core, typically surrounded by Cu(I)-thiolate motifs and the protein shell. We hypothesize that NADH-mediated reduction under physiological conditions enables controlled formation of a Cu^0^ core while preserving Cu(I)-thiolate motifs and protein conformation, thereby governing the optical and biological functionality of the nanoclusters.

Most remarkably, the Pap–CuNCs, when conjugated with anticancer drug 5-fluorouracil, show a significant (∼50-fold) decrease in the half-maximal inhibitory concentration of the drug in HeLa cells, while being non-toxic to normal cells as demonstrated by MTT-based viability assays. Additionally, the system exhibits secondary functionalities such as selective Fe^3+^ sensing and temperature-responsive emission behavior. Molecular dynamics and dynamic docking studies were employed to gain molecular-level insight into the interplay between nanocluster structure and protein conformation.

## 2. Experimental Section

### 2.1. Materials

Papain was purchased from HIMEDIA. Nicotinamide adenine dinucleotide reduced (NADH, disodium salt), 1-ethyl-3-(3-dimethyl) carbodiimide (EDC), 5-fluorouracil (5-FU), Amicon Ultra Centrifugal Filter and TEM Grid were purchased from Sigma. Copper (II) sulfate (CuSO4) and Sodium Hydroxide (NaOH) Pellets were purchased from SRL. To examine the metal ion sensing the following salts of different metal ions were purchased either from SRL or Avra Chemical: Ferrous Chloride (FeCl_2_), Ferric Chloride (FeCl_3_), Stannous Chloride Dihydrate (SnCl_2_.2H_2_O), Lead (II) Sulphate (PbSO_4_) Manganese Chloride (MnCl_2_), Magnesium Chloride (MgCl_2_), Nickel Sulphate (NiSO_4_), Cobalt Nitrate Hexahydrate [Co(NO_3_)_2_.6H_2_O], Cadmium Chloride Monohydrate (CdCl_2_, H_2_O), Mercuric Chloride (HgCl_2_), Barium Chloride Dihydrate(BaCl_2_.2H_2_O), Calcium Chloride (CaCl_2_), Zinc Sulphate Heptahydrate(ZnSO4.7H2O) Potassium Chloride (KCl) and Sodium Chloride (NaCl).

### 2.2 Biocompatible Synthesis of Papain-coated Copper Nanocluster (Pap-CuNC) at Ambient Temperature and Physiological pH

An appropriate amount of papain solution (10 mg/ml, 3 ml) is mixed with an aqueous solution of Copper (II) Sulphate (1.4 mM, 1.5 ml). The pH of the reaction mixture was maintained at physiological pH (∼7) by adding a few drops of NaOH. Then, an appropriate quantity of biological reducing agent (NADH, 10 mg/ml, 1.2 ml) was added dropwise to the mixture under constant stirring at room temperature. The reaction mixture was kept overnight under constant stirring at room temperature. Following the completion of the reaction, the precipitate was collected by centrifugation (at 5000 rpm for 5 mins), and it was resuspended in grade 1 water and washed thrice with grade 1 water using the sedimentation and resuspension method. After that, the solution was stored at 4 °C under dark conditions for further use.

### 2.3 Bioconjugation of Pap-CuNC with 5-fluorouracil (5-FU)

The above-synthesized papain-coated copper nanocluster (Pap-CuNC) were bio conjugated with the anti-cancer drug, 5-FU using simple modification of the previously reported procedure for coupling of the carboxylate group of papain with 5-FU^[28]^ using the coupling agent 1-ethyl-3-(3-dimethyl) carbodiimide (EDC). For that, 1 mg of 5-FU was mixed with 1 mg of Pap-CuNC in a 50 mM HEPES buffer solution at a temperature of 30 ◦C. The EDC solution (5 mM) was added in aliquots in the above solution within 4 hours. After completion of the reaction the drug-loaded Pap-CuNC was initially precipitated by centrifugation (at 5000 rpm for 5 mins) and then purified using an Amicon Ultra Centrifugal Filter (cutoff 30kD). The percentage of drug (5-FU) loading of 5-FU on Pap-CuNC was calculated using the following equation^[28]^

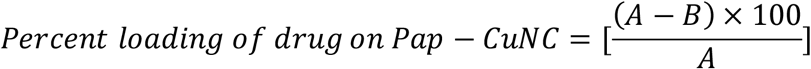

Where A represents the characteristic absorbance of total 5-FU at 265 nm at the beginning of the reaction and B stands for the characteristic absorbance of remaining 5-FU at 265 nm in the supernatant at the end of the reaction. The percentage loading of the drug on Pap-CuNC was found to be around 71%.

### 2.4. Characterization

All fluorescence experiments were performed using a HITACHI fluorescence spectrophotometer, model F-2700. Experiments were carried out in a transparent quartz cell measuring 1.0 cm. A circulating water bath is connected to the instrument for temperature-dependent study. All UV-Vis studies were carried out at room temperature using a Shimadzu twin beam UV-Vis spectrophotometer (model 1800). Circular dichroism (CD) spectra were measured using a Jasco J-810 instrument. The Fourier transform infrared spectroscopy measurements were carried out PerkinElmer Spectrum Two FTIR Spectrometer. The X-ray Photoelectron Spectroscopy (XPS) measurement was carried out employing the AXIS SUPRA system using a monochromatic Al X-ray source. The dynamic light scattering experiments were carried out using Litesizer DLS 500 instruments. The transmission electron microscopy (TEM) measurement was carried out using JEOL JEM-F200 equipped with field emission gun operating at a voltage of 200 kV. The Pap-CuNC dispersion was drop-cast onto carbon-coated Cu TEM grids and then dried overnight under ambient conditions in a vacuum. A time-correlated single photon counting (TCSPC) setup by Horiba (Fluoromax-4 spectrofluorometer) was employed to record the fluorescence lifetime with 340 nm excitation.

### 2.5. Cell Culture

Human embryonic kidney cells (HEK, Normal) cells and HELA (cancer) cells were grown in DMEM (Dulbecco’s Modified Eagle Medium) with 10% fetal bovine serum and 1% Pen Strep Glutamine (from Gibco) in an atmosphere of 5% (v/v) CO_2_ enriched air at 37 ◦C.

### 2.6. Cell Viability Study

The cellular viability of the nanocluster was assessed using MTT assay. Briefly, the cells will be incubated with nanoclusters at a variety of concentrations in a 96-well microplate for 24 h. Post incubation 10 μL of MTT (50 μg/well) was added to each well and incubated for 2-4 hours at 37°C in a humidified atmosphere with 5% CO2. DMSO was used to dissolve the formazan crystals and absorbance at 570 nm was recorded using a microplate reader.

### 2.7. Ion and Temperature Sensing Study

The emission intensity of the nanocluster was measured in varying temperatures, and metal ion type, and concentration from where the ion and temperature sensing properties of the nanocluster is evaluated.

### 2.8. Molecular Dynamics (MD) Simulation of free Papain

A high-resolution crystal structure of Papain has been collected from protein data bank having the PDB ID 1PPP.pdb.^[32]^ We first simulate the free Papain without any bound ligand. We have performed all-atom MD simulation of Papain by using GROMACS (version 2024)^[33]^, and CHARMM36 force field^[34]^ was used for these simulations. The system was first subjected to an energy minimization by using the steepest descent algorithm. Then, this energy-minimized structures were solvated by TIP3P water model^[35]^ in a cubic box. In all the simulations, we used periodic boundary conditions. An appropriate number of counterions was added to make the solvated protein charge-neutral followed by an energy minimization of the solvated system. Then, it was subjected to 1 ns of position-restrained dynamics using an NPT ensemble, by applying a restraining force of 2.5 kcal mol^-1^ Å^-2^ for 1 ns to the protein, whereas all the surrounding water molecules were allowed to move independently. Following the position-restraint simulations, we executed the actual production simulations. We have utilized the particle mesh Ewald method to handle electrostatic interactions. The temperature and pressure of those systems were maintained in an NPT ensemble. A time step of 0.001 ps was used for all these simulations. We performed 20 ns of production simulations and then calculated the helix properties from the simulation trajectories.

### 2.9. Molecular Dynamics (MD) Simulation of Papain-CuNC complex

To prepare the Pap-CuNC composite structure, we first created a stable copper nanocluster (CuNC) using the nanomaterial builder in CHARMM^[36,37]^. The prepared Cu_19_ nanocluster was then subjected to incorporate in a cubic water box and simulated for 10 ns in an NVT ensemble. After the simulation, the stabilized CuNC was allowed to incorporate to the free Papain by dynamical docking approach. Cu19 cluster has increased stability^[38–40]^ (Supporting Video 1). The Cu19 cluster is seen to be stabilized noncovalently in the vicinity of the active site of Papain, in the neighborhood of cysteine residues, notably Cys-25, and in association with polar and aromatic amino acids. Next, the whole system in solvated in a cubic water box and then we performed energy minimization followed by position restrained simulation using GROMACS (version 2024), and CHARMM36 force field was utilized for the simulations using similar protocol as we discussed in previous section. We then performed 10 ns of steer molecular dynamics simulation to keep the nanocluster bound with Papain using a force constant of 5 kcal mol^-1^ A^-2^. After this, we performed 20 ns unrestrained final production simulations with the nanocluster bound Papain.

### 2.10. Docking and MD simulation of 5-FU bound Pap-CuNC

To understand the effect of 5-FU drug binding to Papain-CuNC complex, we first incorporated the drug in the Papain-CuNC composite system. The docking of 5-FU to the Papain system was carried out using the standard settings of Autodock^[41]^ and the lowest energy most favorable binding modes were considered for 5-FU bound to papain among several bound states. The criteria for the selection of the most likely pose of the binding was the most favorable binding energy as calculated by the Autodock mapping results. During the process of docking, our systems were flexible. We found that the most favorable binding pose of 5-FU is associated with its binding near the Papain nanocluster and also the close to a key active site Asp-158 residue in Papain. For MD simulation, the parameters of 5-FU were generated by the CHARMM General Force Field (CGENFF) tool^[42]^. Next, with the 5-FU bound Pap-CuNC system, we performed explicit solvation and then energy minimization, followed by equilibration, and final unrestrained production simulation in the same way as described in previous section.

## 3. Results and discussions

### 3.1. Photophysical characterization of Pap-CuNC

The optical properties of the papain-stabilized copper nanoclusters (Pap-CuNCs) have been studied through the application of UV-visible absorption spectroscopy and photoluminescence spectroscopy. The UV-visible absorption spectrum (blue line, Figure 1) has two distinct bands: a broad feature centered around ∼320 nm (hump 2, Figure 1) and one band centered around ∼249 nm (hump 1, Figure 1A), along with a gradual increase in absorption below 230 nm. The band centered around ∼250 nm, along with the increase in absorption below 230 nm, is attributed to the intrinsic absorption of the papain protein, which occurs through the absorption of aromatic amino acids^[43]^. The feature centered around ∼320 nm might be attributed to ligand-to-metal or metal-ligand charge-transfer transitions, which involve the copper core of the nanoclusters.^[44]^ The absence of a surface plasmon resonance (SPR) band centered around ∼600 nm indicates that larger copper nanoparticles are not formed, thus confirming the formation of ultrasmall nanoclusters^[10]^.

**Figure 1.**
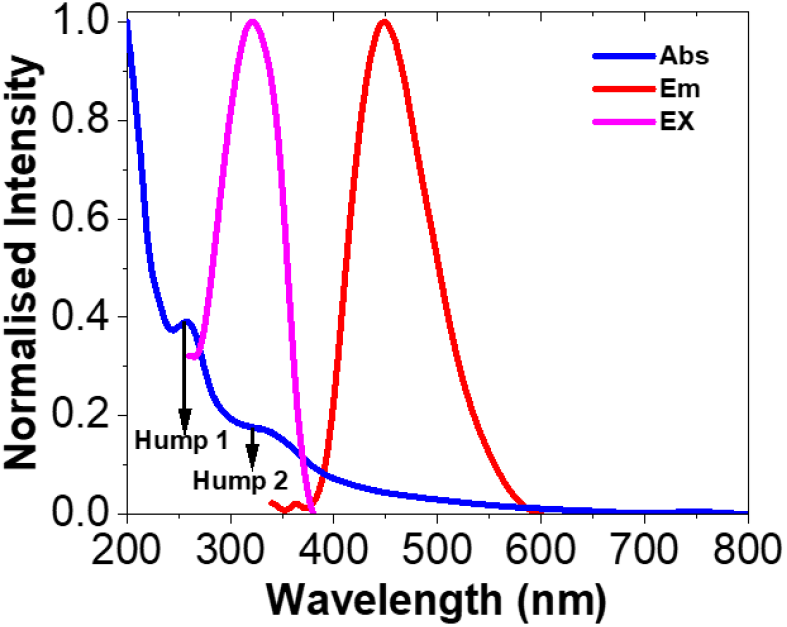
Photophysical characterization of papain-coated copper nanoclusters: UV– visible absorption (blue), excitation (magenta, λ_em_ =449 nm), and emission (red, λ_ex_ =320 nm) spectra of Pap–CuNCs showing a characteristic emission at 449 nm upon excitation at 320 nm, consistent with the formation of luminescent copper nanoclusters.

Upon excitation at 320 nm, Pap–CuNCs show intense blue emission at 449 nm along with a substantial quantum yield of 0.18 (red line, Figure 1). The emission maxima of the Papain and NADH solutions, in the absence of Pap–CuNCs (Figure S1A), were found to be at 370 nm and 497 nm, respectively, with much lower intensity. The difference in emission maxima and intensity of Pap–CuNCs compared to the individual components confirms that the observed photoluminescence is due to the Pap–CuNCs.

The emission maxima of Pap–CuNCs remain nearly constant even after changing the excitation wavelength from 280 nm to 380 nm, which is within ±3 nm (Figure S1B), indicating a nearly uniform distribution of emissive states in the Pap–CuNCs. On the other hand, the emission intensity of Pap–CuNCs depends significantly on the excitation wavelength, and the maximum intensity is observed at 320 nm (Figure S1C), which is in accordance with the maximum in the excitation spectrum (magenta line, Figure 1) when λ_em_ is fixed at 449 nm, indicating that the emission is due to absorption at 320 nm. Furthermore, the Pap–CuNCs exhibit excellent photostability, retaining their emission intensity over extended storage (∼1 month, Figure S1D), highlighting their suitability for long-term bioimaging and sensing applications.

### 3.2. Characterization of Protein Corona using Circular-Dichroism (CD) and Fourier Transform Infrared (FTIR) Spectroscopy

CD spectroscopic study (Figure 2A) reveals an increase in the 222 nm band while a decrease in the 208 nm band in the CD spectrum of Pap–CuNCs compared to native papain. Using the ChiraKit software analysis^[45]^, we observed an increase in the α-helical content of protein in Pap-CuNCs to ∼31% from 26% in native papain, whereas the β-sheet content decreases to 19% from 27%, respectively. Altogether, the CD spectrum indicates partial restructuring of the protein secondary structure, suggesting interaction of copper nanoclusters with the protein backbone leading to increased α-helical propensity. Notably, this experimentally observed increase in α-helical content is consistent with molecular dynamics simulations, which reveal stabilization of the protein structure upon nanocluster binding (section 3.5.).

**Figure 2.**
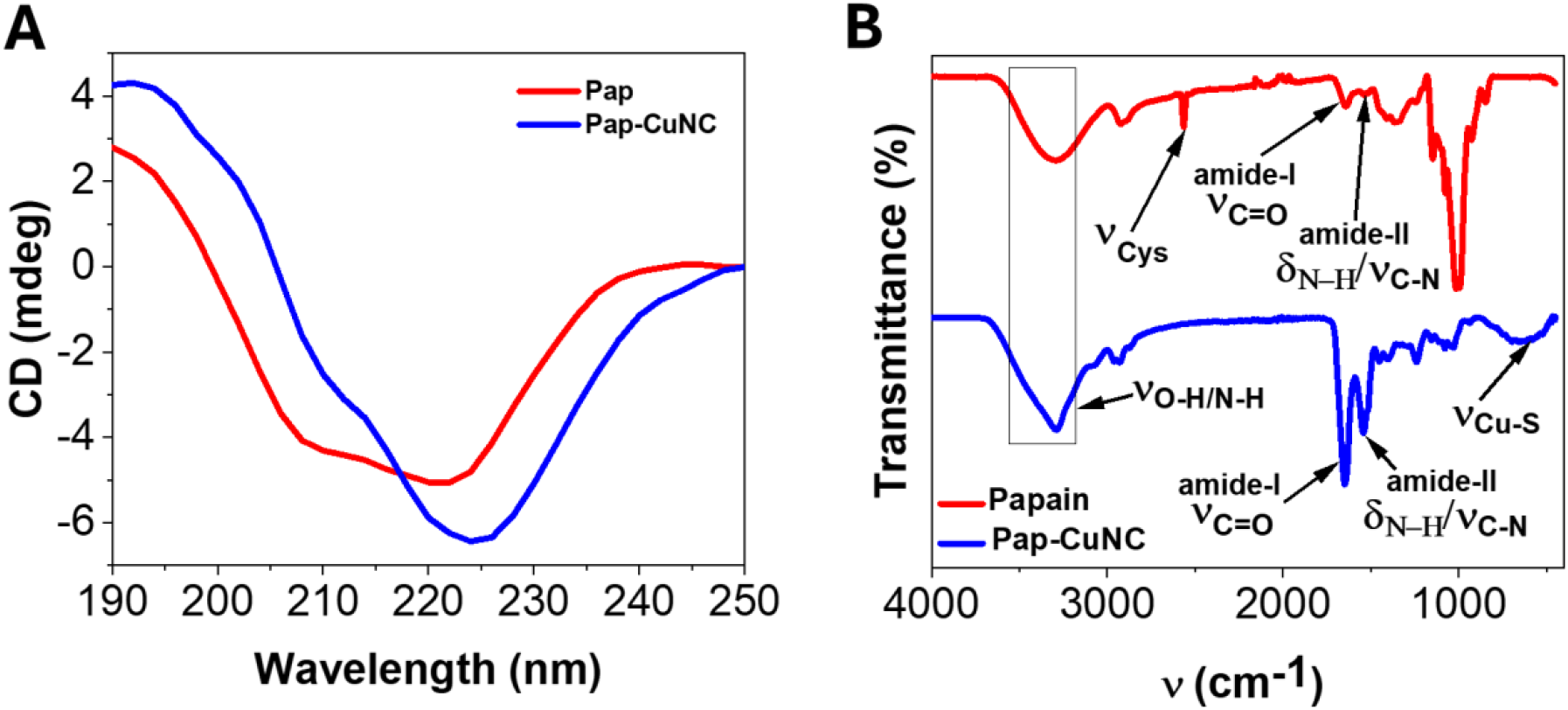
Characterization of Protein Corona of Pap-CuNC: A. CD spectra of Papain (red) and Pap-CuNC (red), indicating the increase in α-helical content of the papain. B. FTIR spectra of Papain (red) and Pap-CuNC (blue).

To further investigate the interaction pattern of protein corona with metallic core, FTIR spectra of papain and Pap-CuNC samples were recorded (Figure 2B). The broad peak observed in the high-wavenumber FTIR spectrum of the protein in the range of 3200-3400 cm^−1^, corresponding to O-H/N-H stretching^[46]^, indicates a slight broadening of the peak upon formation of the Pap-CuNC, indicating the possible involvement of hydroxyl groups in hydrogen bonding with the nanoclusters. The characteristic amide I peak observed in the FTIR spectrum of the protein at ∼1650 cm^−1^, corresponding to C=O stretching, and the amide II peak observed at ∼1540 cm^−1^, corresponding to N-H bending along with C-N stretching^[46]^, indicate subtle changes in intensity and position after nanocluster formation, indicating a subtle change in the protein backbone structure during coordination with copper. In addition, the band around **∼**1000 cm^−1^, attributed to ν(C–O) stretching vibrations of hydroxyl and carboxyl groups in papain^[46]^, is significantly diminished or absent in Pap–CuNC. This disappearance indicates the direct participation of these oxygen-containing groups in copper coordination, leading to the formation of Cu–O linkages and consequent suppression of the C–O vibrational mode. Importantly, the thiol group stretching of cysteine residues observed as a weak peak near ∼2550 cm^−1^ indicates^[47]^ a significant reduction in peak intensity upon formation of the Pap-CuNC, indicating the possible coordination of thiol groups with metal coordination. The broad peak observed near ∼600-620 cm^−1^ in the FTIR spectrum of the Pap-CuNC, corresponding to the Cu-S stretching,^[48,49]^ while this feature is absent in the spectrum of native papain. Altogether, those spectral features suggest the stabilization and coordination of copper atoms by the thiolate groups of cysteine residues, along with amide and hydroxyl functionalities of the protein, during the formation of Pap-CuNC.

### 3.3. Characterization of Metal Core using X-ray Photoelectron Spectroscopy (XPS), Dynamic Light Scattering (DLS) and High-Resolution Transmission Electron Microscopy (HR-TEM) studies

Cu has the lowest reduction potential of the three coinage metals, making it quickly oxidize. As a result, it was judged necessary to determine the oxidation state of Cu in the composite. For this, XPS analysis (Figure 3 and S2) was used to confirm the oxidation state of Cu in the Pap-CuNC (Figure 3A). Two significant peaks at ∼951.9 EV and ∼932.4 eV were ascribed to Cu ^2^p_1/2_ and Cu ^2^p_3/2_, which were indicative of Cu^0^.^[50]^ (Figure 3A) The absence of a Cu^2+^ satellite peak at around 940 eV suggests that the Pap-CuNC did not contain Cu^2+^. However, given that the signal owing to Cu^+^ appears at 0.1 eV away from Cu^0^, its presence cannot be conclusively ruled out. The O 1s (Figure S2D) peak consists of two prominent peaks at ∼531.6 eV (C=O) and ∼533 eV (C-O/C-OH),^[51–53]^ which indicate the involvement of the oxygen groups in the papain molecule in weak coordination with the Cu surface. In the S2p spectrum (Figure S2B), the prominent peak at ∼162.8 eV corresponds to the thiolate bonds in the cysteine residues.^[54]^ Therefore, it is clear that the anchoring is done by the sulfur atoms, which provide strong attachment to the Cu nanocluster, whereas the secondary interactions are provided by the oxygen atoms, thereby forming a stable protein corona with the Cu core.^[50]^

**Figure 3.**
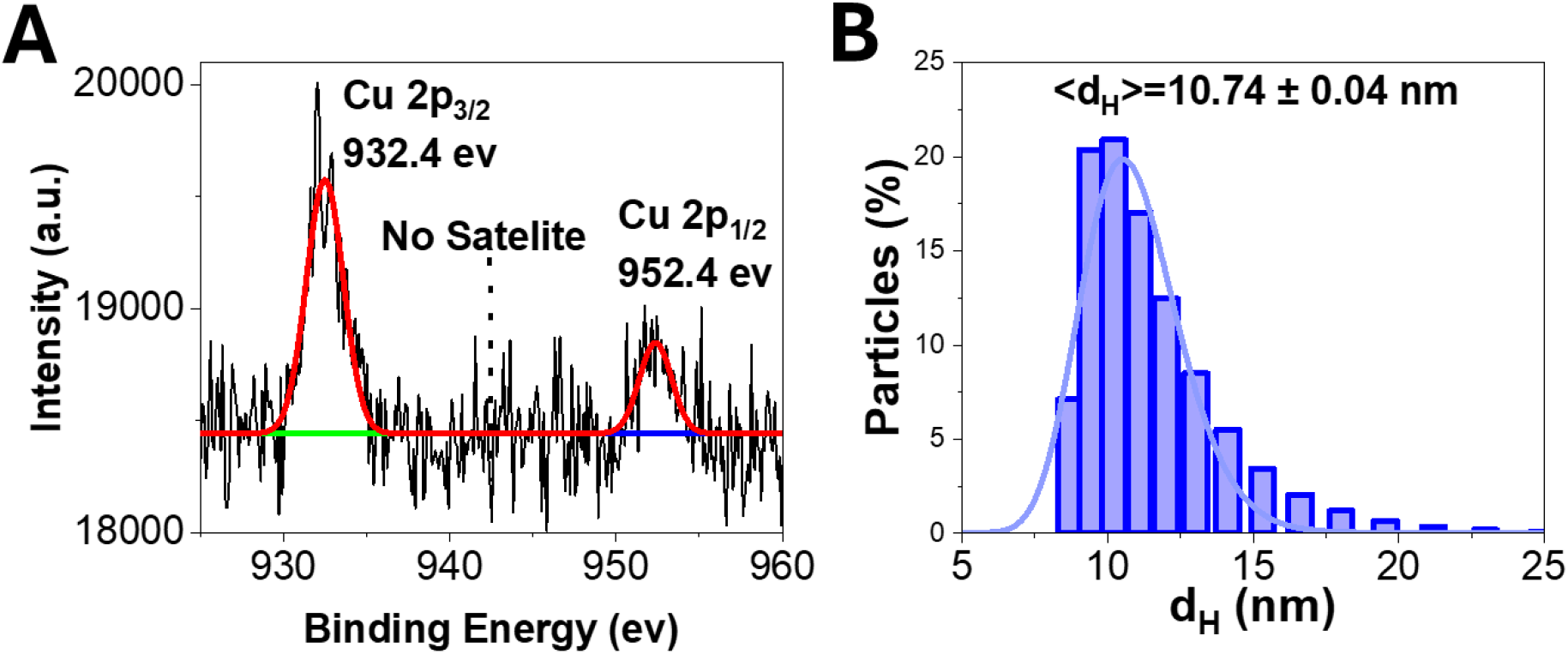
A. XPS spectrum of Cu 2p electrons in Pap-CuNCs. The absence of a satellite peak is indicative of the absence of Cu^2+^ species. B. Distribution of hydrodynamic diameter (d_H_) Pap-CuNC as measured by Dynamic Light Scattering (DLS).

The size and structural properties of the Pap-CuNC were investigated using DLS (Figure 3B) spectroscopy and HR-TEM images (Figure 4). DLS analysis shows that the hydrodynamic diameter of the Pap–CuNCs is centered at ∼10.74 ± 0.04 nm, which indicates the formation of a protein corona. The Pap-CuNC exhibits a mean zeta potential of −12.39 mV. This shows that Pap-CuNC has a moderately negative charge due to deprotonated carboxyl groups (-COO-) and hydroxyl groups of papain, indicating that the colloidal solution is moderately stable. This stability is due to electrostatic repulsion and steric hindrance. This agrees with the XPS and FTIR results showing Cu–S and oxygen-based interactions between the protein corona and the Pap-CuNC. This slightly negative charge is suitable for biological purposes as it increases dispersibility while reducing nonspecific interactions.

**Figure 4.**
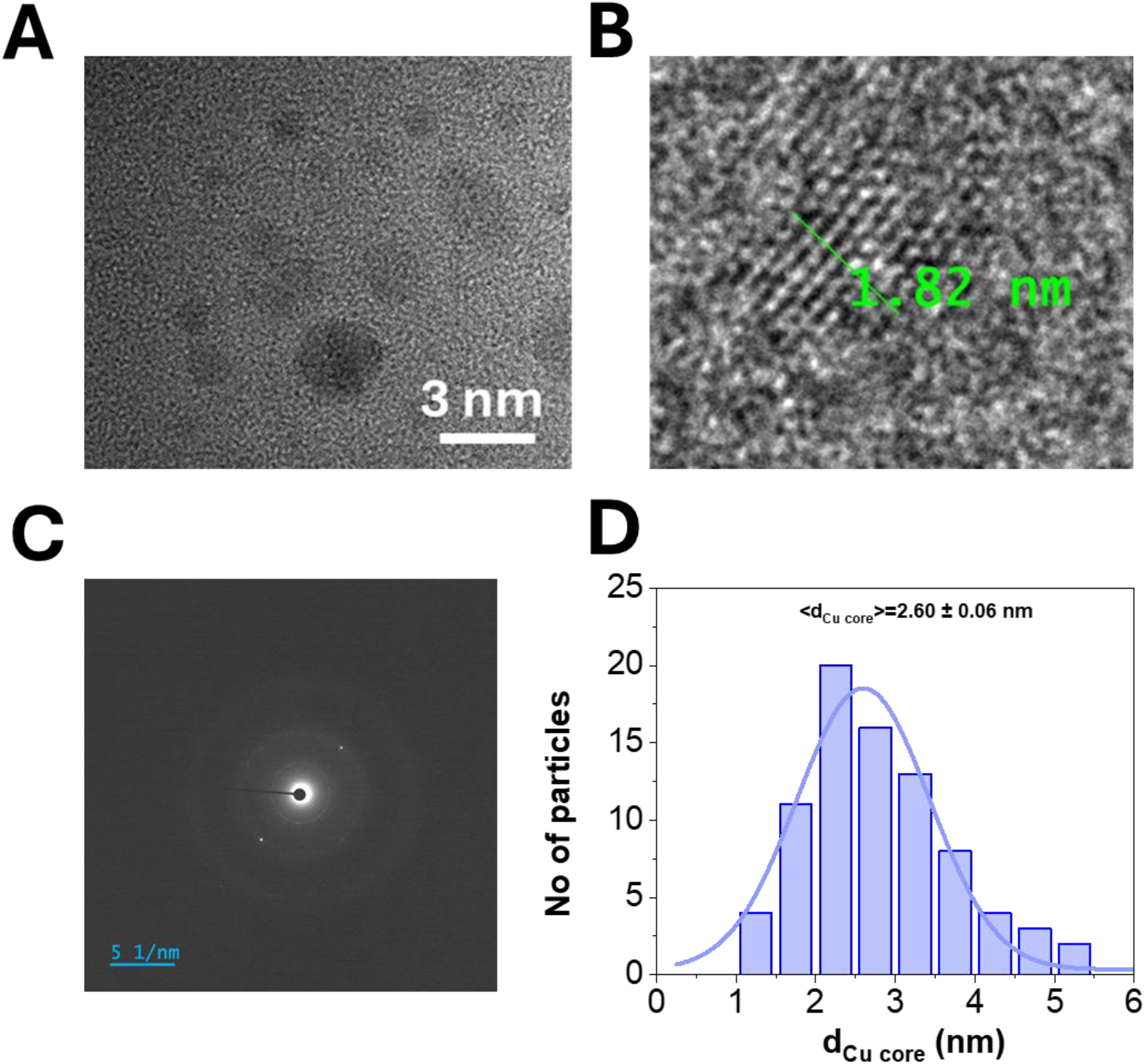
Characterization of Metallic Core of Pap-CuNC: A. HRTEM image of Pap-CuNC showing the distribution of metallic core of the nanocluster B. Zoomed image of a single particle showing the crystal planes. Interplanar spacing ∼ 0.2065 nm consistent with a nanocrystalline FCC Cu^0^ core, as further supported by lattice analysis (Figure S3) C. Selected Area Electron Diffraction (SAED) pattern showing nanocrystalline domains embedded within a partially disordered matrix consistent with nanocrystalline or polycrystalline behavior. D. Distribution of the size of the metallic core of Pap-CuNC as measured from HRTEM images.

On the other hand, TEM images (Figure 4 A-C) demonstrate well-dispersed nanocrystalline metal nanoclusters of smaller core sizes. Statistical analysis of the obtained results indicates that the average value of the core diameter of the nanoparticles is ∼2.60 ± 0.06 nm (Figure 4D), consistent with the formation of ultrasmall nanoclusters. Clear lattice fringes in the HR-TEM images (Figure 4 B) confirm the crystallinity of the metal core of the nanoparticles. Selected Area Electron Diffraction (SAED, Figure 4C) pattern demonstrates nanocrystalline domains embedded within a partially disordered matrix consistent with the characteristics of polycrystalline or nanocrystalline materials. Fast Fourier Transform analysis and filtering using the inverse FFT algorithm helped determine the interplanar distance of the crystal lattice (Figure S3A). By using the line profile of the fringes, the interplanar spacing is found to be ∼0.2065 nm, corresponding to the (111) plane of face-centered cubic crystalline copper.^[50]^ Clear diffraction spots in the FFT image supports the crystalline nature of the Cu^0^ core. The apparent difference between the hydrodynamic diameter and the core size clearly indicates a core–shell architecture, where the metallic Cu nanocluster core is encapsulated within a protein matrix. Energy Dispersive X-ray Spectroscopy (EDS) confirms the elemental composition of the Pap–CuNCs, showing characteristic Cu peaks (Cu L and K lines) along with C, N, and O signals (Figure S3B) from the papain matrix, thereby verifying the formation of a copper nanocluster core stabilized by a protein corona. Overall, these results confirm the successful formation of crystalline, ultrasmall copper nanoclusters stabilized by papain under biocompatible conditions.

In summary, from a mechanistic point of view, the observed blue emission is ascribed to the quantum confinement of the electronic structure of the copper core, influenced by the Cu(I)-thiolate motifs on the surface and the protein corona effect. The larger size of the metal core from the TEM analysis indicates that the observed emission is from localized electronic states in a metal core rather than a molecule-like structure. The surrounding proteins would stabilize these electronic states and suppress non-radiative decay pathways.

### 3.4. Application of Pap-CuNC

#### A. Major Application: Very proficient cancer drug carrier

To test the in vitro toxicity of Pap-CuNC, we have performed MTT based cell viability assay (Figure 5). The viability of HEK cells after incubating the cells with the Pap-CuNC for 24 hours was studied as a function of the concentration of the nanocluster. The Pap-CuNCs demonstrated remarkable biocompatibility (∼ 80% viability) even at very high concentrations (∼4000 μg/ml) (Figure 5A), clearly depicting the non-cytotoxicity of the nanocluster, confirming that the Pap-CuNC is suitable for intracellular studies. However, in the case of HeLa cells, the Pap-CuNCs show higher toxicity with an IC50 value of 296.9±51.8 μg/ml (Figure 5B). It has been reported earlier that papain causes apoptosis of cancer cells while being nontoxic to normal cells^[55,56]^. Thus, the papain coating of the copper nanocluster on Pap-CuNCs may be responsible for its toxicity towards cancerous HeLa cells while being non-toxic to normal HEK cells. Then we covalently attached cancer drug Fluorouracil (5-FU) with the Pap-CuNCs and performed MTT-based cell viability assay of HeLa cells in presence of only 5-FU (Figure 5C) and 5-FU-attached Pap-CuNCs (Figure 5D). We observed the IC50 value of 5-FU decreases drastically from 170.3±31.2 μg/ml (note, previous literature reported slightly higher IC_50_ value of 5-FU in HeLa cell ∼ 251 μg/ml ^[57]^) to 3.3±1.2 μg/ml in the case of 5-FU attached Pap-CuNCs. Thus, Pap-CuNCs demonstrates strong potential as an efficient cancer drug carrier. This enhanced therapeutic performance is consistent with the nanocluster-induced stabilization of protein structure observed in simulation and spectroscopy.

**Figure 5.**
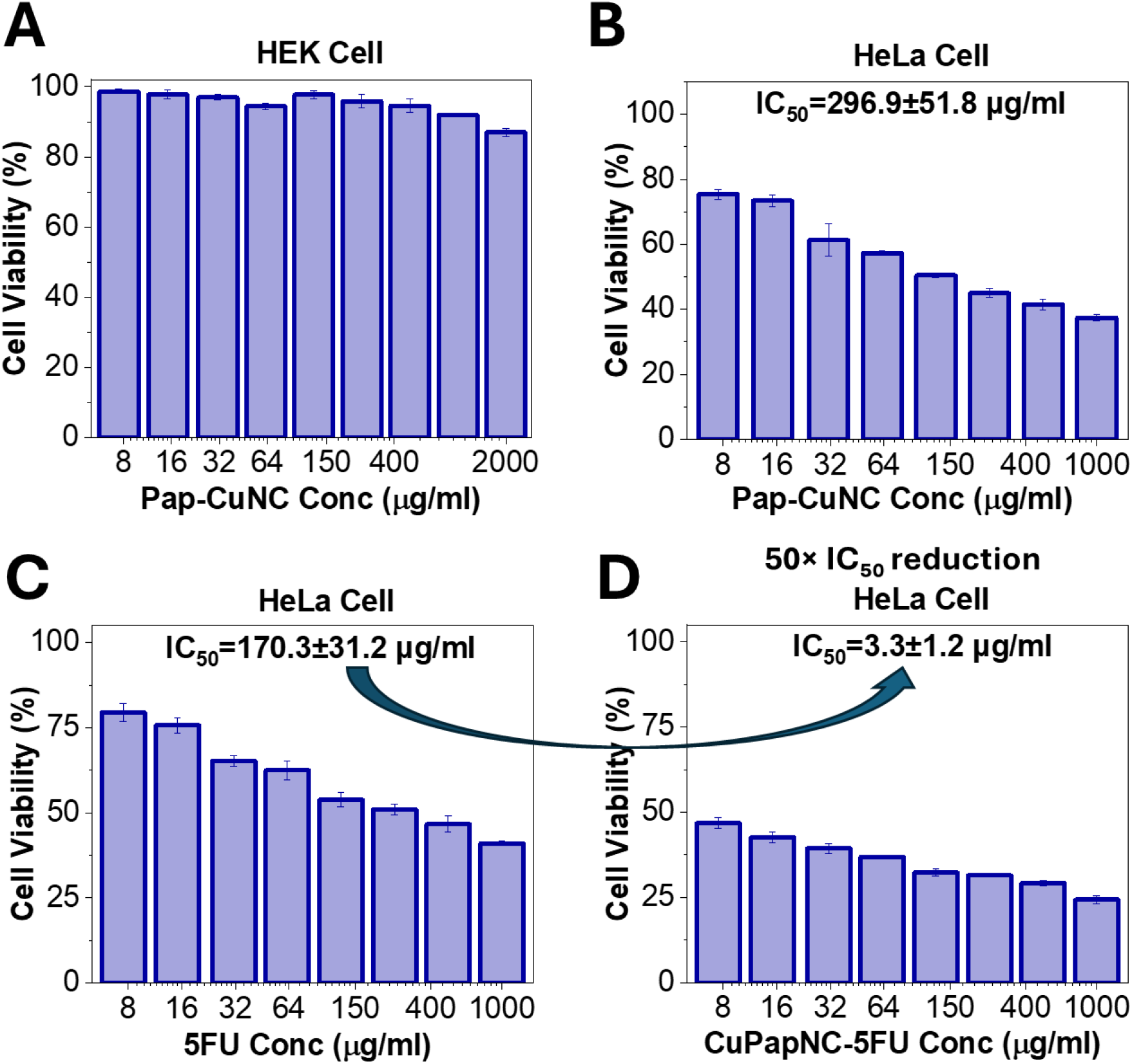
Pap–CuNCs as an efficient nanocarrier for anticancer drug delivery. MTT-based cell viability assays showing cytotoxicity profiles of (A) Pap–CuNCs toward HEK (normal) cells, demonstrating high biocompatibility, and (B) Pap–CuNCs toward HeLa (cancer) cells. (C) Cytotoxicity of free 5-fluorouracil (5-FU) in HeLa cells, and (D) enhanced cytotoxicity of 5-FU-conjugated Pap–CuNCs in HeLa cells. The nanocluster– drug conjugate exhibits a dramatic reduction in IC50 (∼50-fold) compared to free 5-FU, highlighting the effectiveness of Pap–CuNCs as a drug delivery platform.

#### B. Secondary Functionalities: Selective detection of Ferric ions and Temperature Sensing Properties of Pap-CuNC

To investigate the metal ion sensing capability of Pap-CuNC nanocluster, it was incubated with appropriate metal ion concentrations for ∼3 mins prior to measurement. An intriguing aspect of the Pap-CuNCs nanocluster is that the PL intensity almost remains constant in the presence of several metal ions (Figure 6A). Iron in the Fe^3+^ state almost completely reduces the PL intensity of Pap-CuNCs nanoclusters and causes visible precipitate development. However, iron in the Fe^2+^ state reduces the PL intensity of Pap-CuNCs nanoclusters by less than 30% without forming precipitates. Thus, this nanocluster may be ideal for identifying the Fe^3+^ concentration for water quality monitoring. The Stern-Volmer plot [F_0_/F versus the concentration of the quencher (Fe^3+^), where F_0_ is the fluorescence intensity without quencher, and F is the intensity with quencher] showed a parabolic pattern (Figure S4A) which can be fitted with a second-order polynomial function. This suggests that the quenching of the fluorescence intensity by Fe^3+^ may be ascribed to combining static and dynamic quenching.^[58]^ This was further supported by liner dependence of (τ_0_/τ) with nanocluster concentrations (Figure S4B). The probable mechanism is represented with a schematic in the supporting information (Figure S4C).

**Figure 6.**
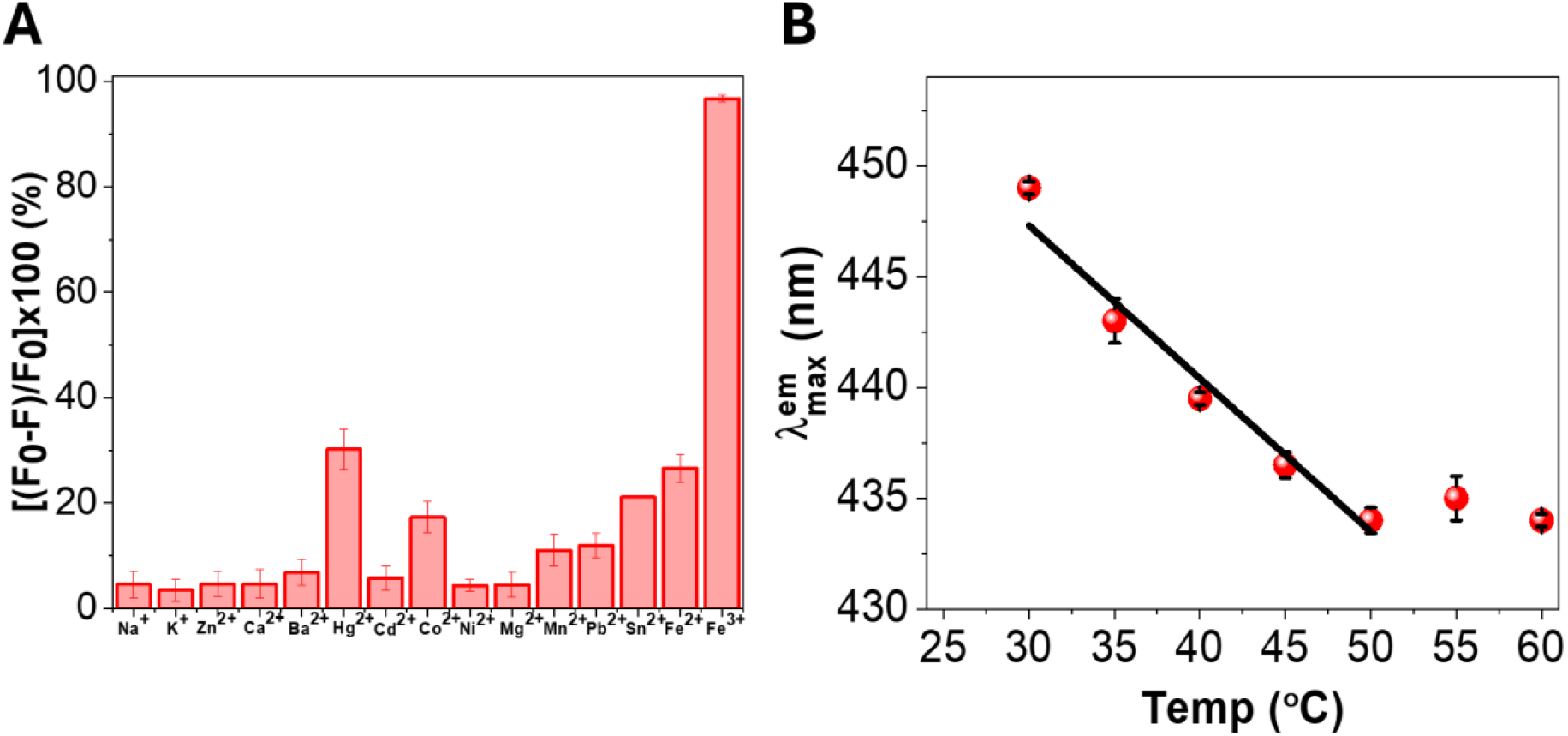
Selective detection of Ferric ions and Temperature Sensing Properties of Pap-CuNC: A. Quenching (%) of Pap-CuNC fluorescence by different metal ions (concentration of ions: 500 μM except Pb^2+^ (92 µM)). F_0_ and F represent the fluorescence intensities in the absence and presence of metal ions, respectively B. Linear change of emission maxima of Pap-CuNC in physiologically relevant temperature range (30-50°C).

The emission spectra of Pap-CuNC show an interesting feature upon temperature variation. Although the emission intensity of the Pap-CuNC remains constant (Figure S5) over a temperature range of 30 ◦C to 60 ◦C range, the emission maxima (λ_max_) decrease linearly (Figure 6B) from 449 nm at 30◦C to 434 nm at 50◦C and then show a plateau up to 60◦C. Although the core temperature of the healthy human body is tightly regulated within a range of 36.5–37.5°C local and intracellular variations create subcellular hotspots where the temperature is 5-10°C higher than the surrounding cytoplasm such, as mitochondria^[59]^. Pathological conditions such as infection, cancer, or inflammatory disease frequently give rise to temperature variability. Determining whether the rate of a patient’s body temperature is changing indicates how their body responds physiologically to the illness/condition.^[60,61]^ Thus, there is a growing interest of development of accurate, high-resolution (in space/time), and sensitive means of acquiring local temperature in different subcellular regions (nano-thermometry). In order to meet this need, fluorescent species agents that respond to temperature by changing their optical properties are of high demand as a suitable candidate for optical nanothermometers.^[60,61]^ The unique property of change of emission maxima on the variation of temperature makes Pap-CuNC a suitable candidate for fluorescent nano-thermometry.

### 3.5. Understanding the Mechanistic Aspect by Dynamic Docking and Molecular Dynamics Simulation study

Jellium Model has been used extensively to theoretically predict the number of metal atoms present in the nanoclusters [E_NC_ = E_f_/(N)^1/ 3^, where E_NC_ stands for the emission energy of the nanoclusters, which corresponds to the emission maxima, E_f_ is the Fermi energy of the metal in its bulk state, and N is the number of atoms in the nanoclusters.]^[8]^ According to this model, papain-coated copper nanocluster has an emission peak of about 449 nm, indicating the presence of Cu_17_ (17 Cu atoms in the metal core) in the NC. However, its use in ligand-protected systems is constrained owing to electron localization and metal-ligand interactions. To generate a realistic structure, molecular dynamics (MD) simulation and dynamic docking approaches have been carried out (Figure 7).

**Figure 7.**
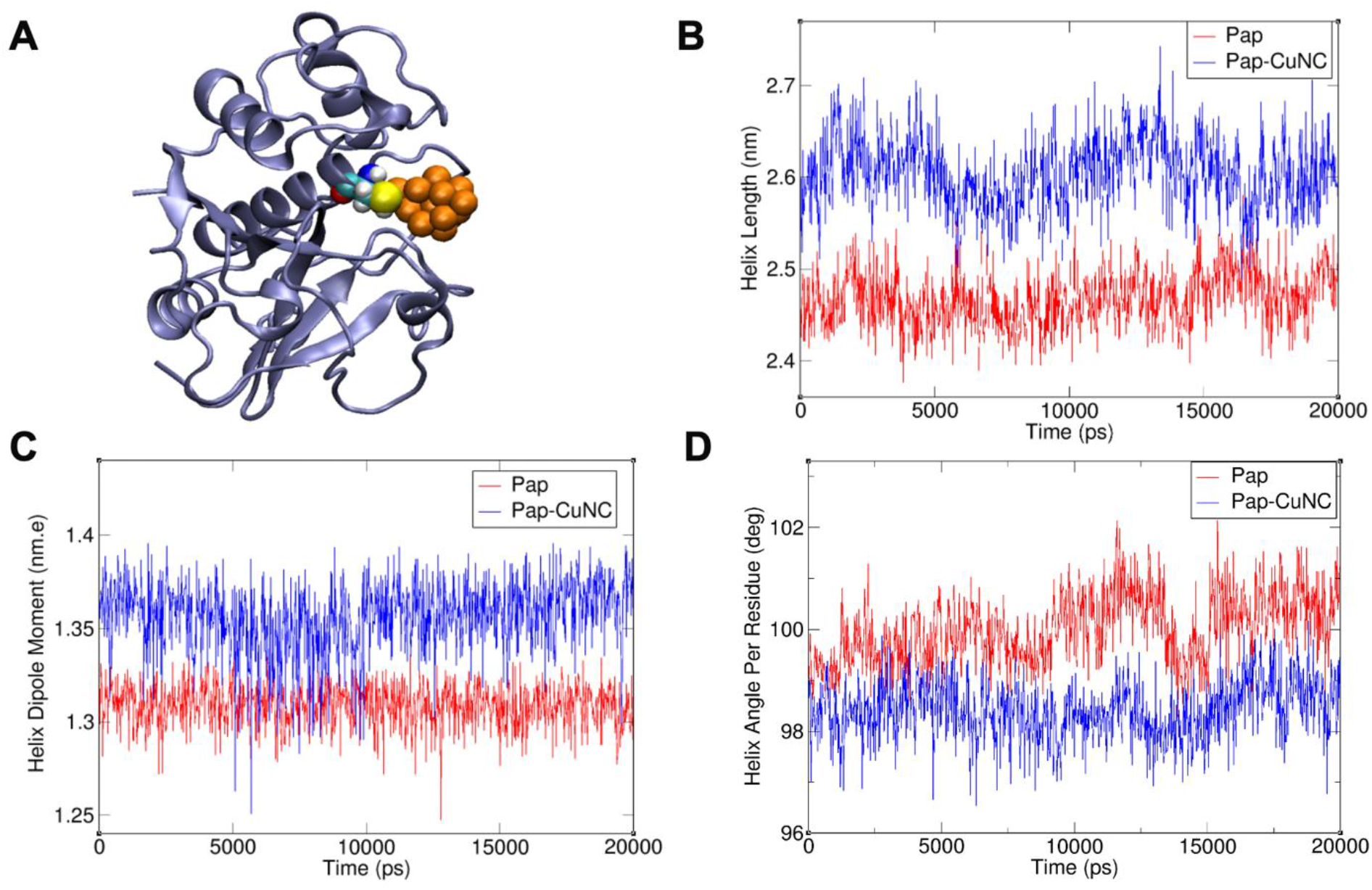
Dynamic Docking and Molecular Dynamics Simulation of Pap-CuNC: A. Docked conformation of Cu_19_ into papain. Papain is rendered in cartoon representation by iceblue color.CuNC is shown by orange color with vdW mode and its binding to a key active site residue CYS-25 is shown, where the CYS-25 is marked by vdW representation. B. Plot of Helix Length vs Time C. Plot of Dipole Moment vs Time D. Plot of Helix Angle per Residue vs Time.

It was observed that the Cu19 cluster (the closest magic number Cu cluster of Cu_17_)^[38–40]^ has increased stability in the presence of papain protein (supporting video 1). The Cu19 cluster is seen to be stabilized noncovalently in the vicinity of the active site, in the neighborhood of cysteine residues, notably Cys-25, and in association with polar and aromatic amino acids (Figure 8). This binding pattern is consistent with the observed Cu-S and Cu-O bond formation from FTIR and XPS studies.

**Figure 8.**
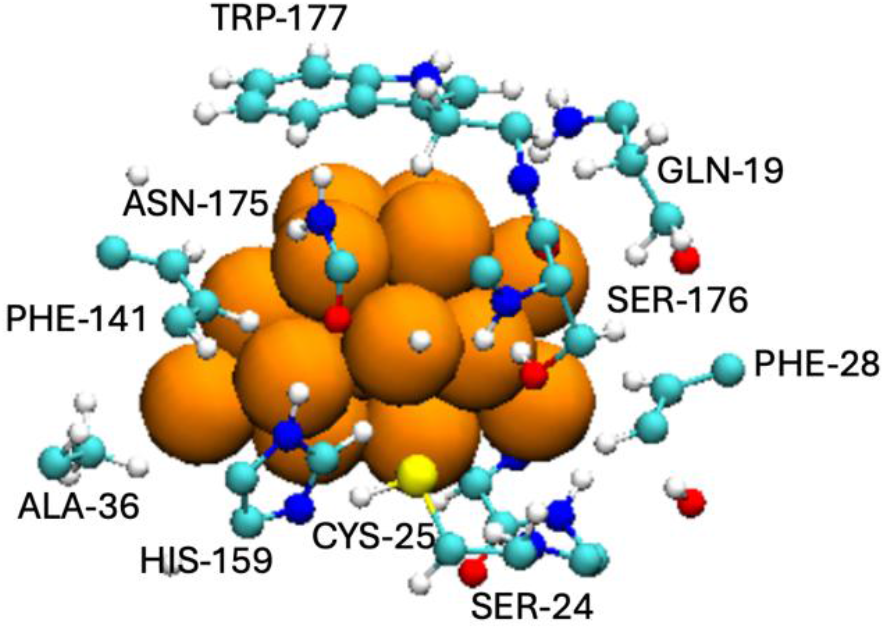
A snapshot of Papain amino acid residues within 0.45 nm of CuNC: The residue identities were marked by their three-letter amino acid code. CuNC is shown by vdW mode in orange color and the amino acid residues were rendered in CPK mode.

It should be mentioned that the Cu19 cluster induces a significant conformational rearrangement of papain. Analysis using MD simulation revealed an increase in helical length (Figure 7B) and helix dipole (Figure 7C), while the helix angle per residue (Figure 7D) was significantly reduced, indicating the stabilization of the α-helix in the protein. All these findings were supported by the results obtained through CD spectroscopy (Figure 2A), which revealed an increase in the α-helix content due to nanocluster formation. Thus, it can be assumed that the copper nanocluster serves as a structural framework, stabilizing the protein conformation.

As the increased stability of bioactive proteins has been shown to enhance their apoptosis capability^[62,63]^, leading to improved therapeutic performance. Therefore, the cytotoxic properties of Pap–CuNCs observed when exposed to cancer cells may partly be attributed to enhanced structural integrity of papain upon nanocluster binding.

Moreover, docking simulations performed on 5-FU (Figure 9) show that the drug is more prone to bind to the surface of the protein close to the nanocluster, possibly via interaction with aspartate residues (ASP-158, Figure 9A) exposed on the protein surface. This prediction is because we have used EDC linkers to couple 5-FU to the carboxylate group of the protein. The high solvent-accessible surface area (SASA) of the bound drug (∼2.5-2.7 nm^2^, Figure 9B) points to considerable solvent which may facilitate efficient drug release. It is important to note that Asp-158 in papain is a crucial residue located near its active site, specifically within hydrogen-bonding distance of the catalytic histidine-159 (His159). Studies indicate it acts as a modulator of the active site environment (electrostatic modulation of the active site residue is triggered by ASP-158 residue of Papain)^[64]^.

**Figure 9.**
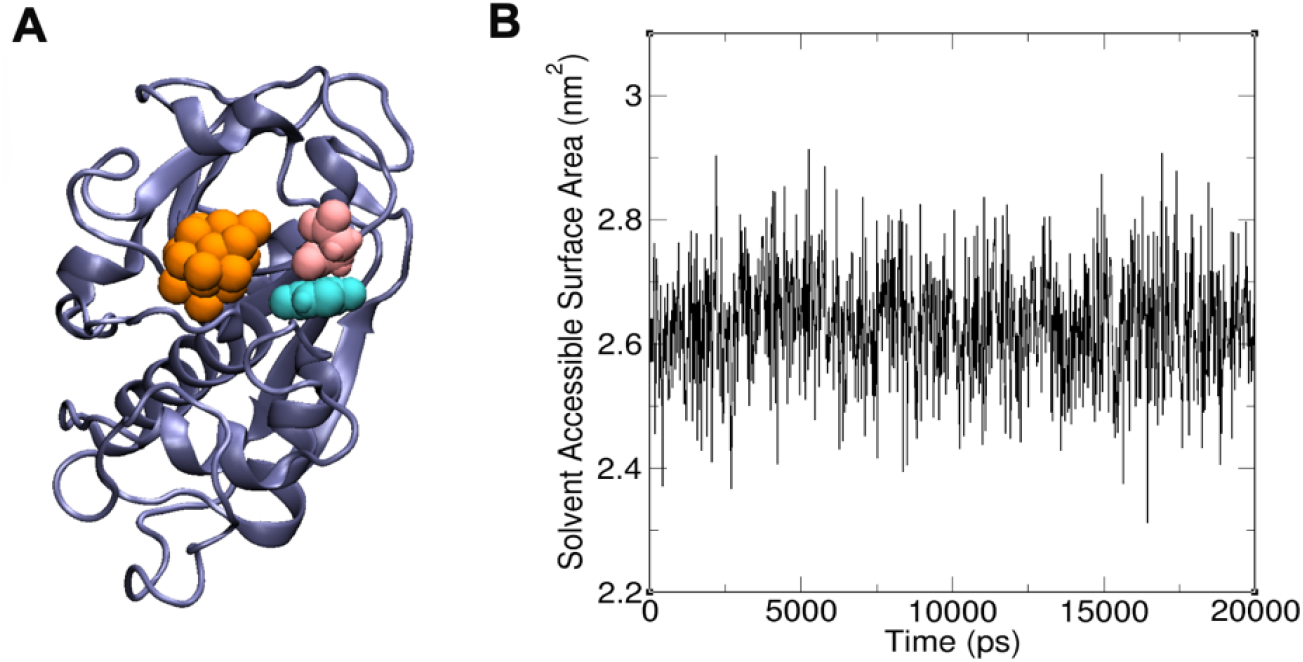
Docking of 5-FU and Molecular Dynamics Simulation Study of binding of 5-FU with Pap-CuNC: A. Docked conformation of 5-FU into Pap-CuNC. The protein is shown in iceblue color, and the Copper nanocluster (CuNC) is rendered in orange color. 5-FU drug and its adjacent interactive ASP-158 is shown by cyan and pink color code respectively. B. Plot of Solvent Accessible Surface Area (SASA) of 5-FU vs Time. Here higher SASA value is indicative of highly solvent exposed nature of 5-FU drug.

In summary, MD and docking simulations have yielded an in-depth molecular analysis of the system, demonstrating the formation of Cu19 to be associated with strong interactions between the protein and the nanocluster, while increased helicity and drug accessibility contribute to superior biological function of the Pap-CuNC construct. These results provide a molecular-level basis for the experimentally observed structure–function relationship, highlighting the role of nanocluster–protein interactions in stabilizing protein conformation.

## 4. Conclusion

In summary, we have employed a physiologically relevant reducing agent, NADH, to develop a biocompatible strategy for the synthesis of protein-stabilized copper nanoclusters. Our approach enables the formation of ultrasmall, luminescent Cu nanoclusters under mild conditions, enhancing the protein’s helicity and function, thereby overcoming a key limitation of conventional synthetic methods. A combination of spectroscopy, microscopy, and computation shows that the nanocluster contains a metallic core composed of Cu^0^ atoms, along with a Cu(I)-thiolate ligand system and a modified protein corona, while MD simulations have shown that a Cu19 cluster is a stable structural scaffold within the protein milieu.

Notably, the NADH-mediated reduction pathway provides control over the electronic structure of the nanocluster, which governs its optical properties and, therefore, its biological behavior. As such, the synthesized Pap–CuNCs have strong photoluminescent characteristics, stability, and multifunctionality such as sensitivity to Fe^3+^ ions as well as thermos responsive emission. Importantly, however, their potential in improving the anticancer effects of 5-fluorouracil makes them a promising drug-delivery system.

Overall, this work demonstrates that the application of redox biology enables a rationale-driven strategy for engineering nano-clusters in a structured manner, thereby opening up new avenues for designing nanomaterials for biomedical and sensing applications. More broadly, this work establishes physiological redox chemistry as a design principle for engineering bio-integrated nanomaterials with controlled structure and function. This strategy is expected to be extendable to other protein–nanocluster systems under biologically relevant conditions.

## Data availability statement

Data and documentation are available upon reasonable request from the corresponding authors.

## Funding statement

This work is supported by the Gandhi Institute of Technology and Management (GITAM): Research Seed Grants (Ref: F.No 2023/0224); Start-up Research Grant (SRG/2023/000147), and Advanced Research Grant (ANRF/ARG/2025/000361/CS) from Anusandhan National Research Foundation (ANRF), Government of India. SG also acknowledges IIT Jammu for providing seed grant support. S.G. acknowledges the Central Instrumentation Facility SAPTARSHI and the High-Performance Computing Facility AGASTYA of IIT Jammu. We acknowledge the use of Circular Dichroism facility of CSIR-Centre for Cellular & Molecular Biology (CCMB). We also acknowledge the use of X-ray Photoelectron Spectroscopy (XPS) and Time-Correlated Single Photon Counting (TCSPC) facility of IIT Hyderabad. A.S. and A.A. acknowledge IIT Jammu for providing fellowship. A.B. acknowledge IIT Hyderabad for providing fellowship. Vanshika is grateful to the University Grants Commission (UGC), New Delhi, for awarding the Junior Research Fellowship (JRF) [Award No.241620182653].

## Conflict of interest disclosure

The authors declare no conflicts of interest.

## Notes

### Competing Interest Statement

The authors have declared no competing interest.

