## Supporting Information for "One-Pot NADH-Mediated Physiological Redox-Controlled Synthesis of Papain-Stabilized Copper Nanoclusters with Preserved Bioactivity for Efficient Drug Delivery"

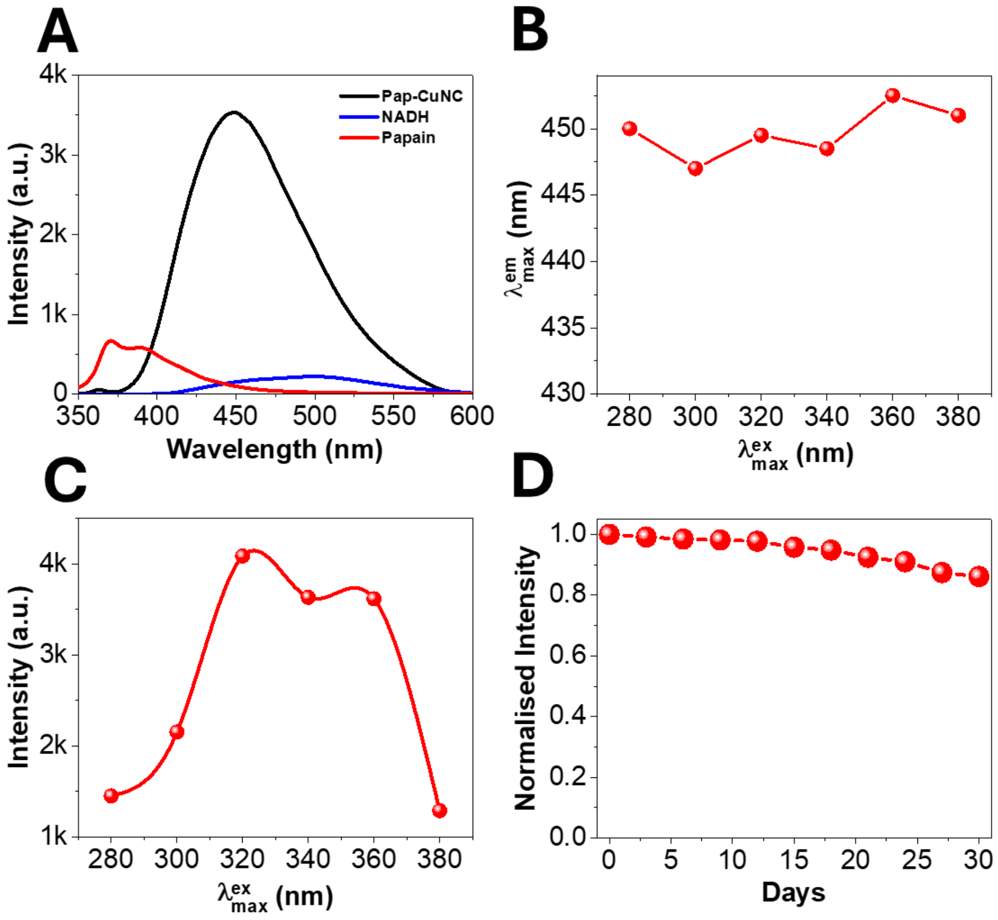


**Figure S1. Photophysical characterization of papain-coated copper nanoclusters:** A. Emission spectra ($\lambda_{ex}$=320 nm) of papain (red), NADH (blue) and Pap-CuNC (black) measured under the same condition ($\lambda_{ex}$=320 nm, pH~7 and concentration (~ 1 mg/ml). B. Dependence of emission maxima ($\lambda_{em}$) of Pap-CuNC on excitation maxima ($\lambda_{ex}$). C. Dependence of intensity at emission maxima ($\lambda_{em}$) of Pap-CuNC on excitation maxima ($\lambda_{ex}$). D. Normalized intensity of Pap-CuNC on excitation maxima ($\lambda_{ex}$) over time.

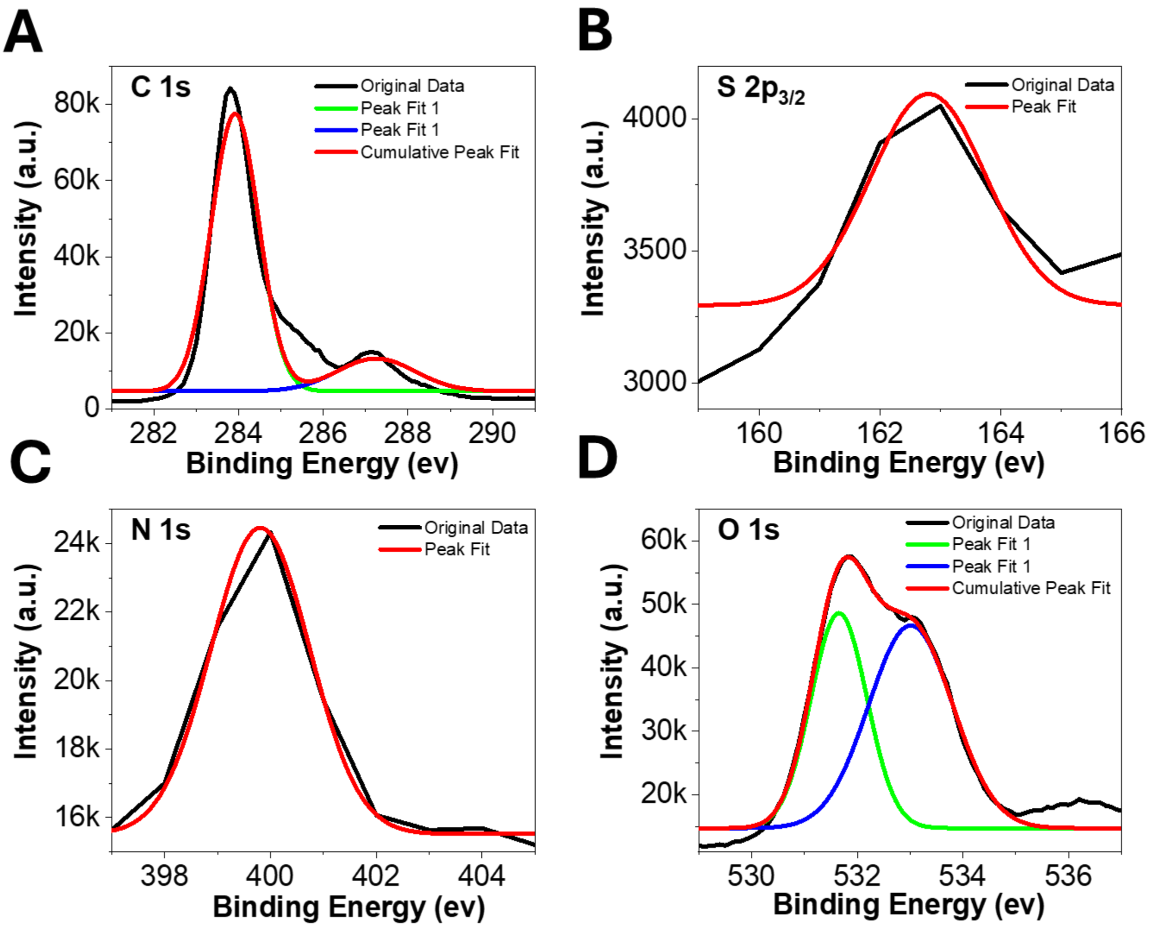


**Figure S2.** X-ray photoelectron spectra of the elements present in the Pap-CuNC. (A) C 1s, (B) S 2p, (C) N 1s and (D) O 1s.

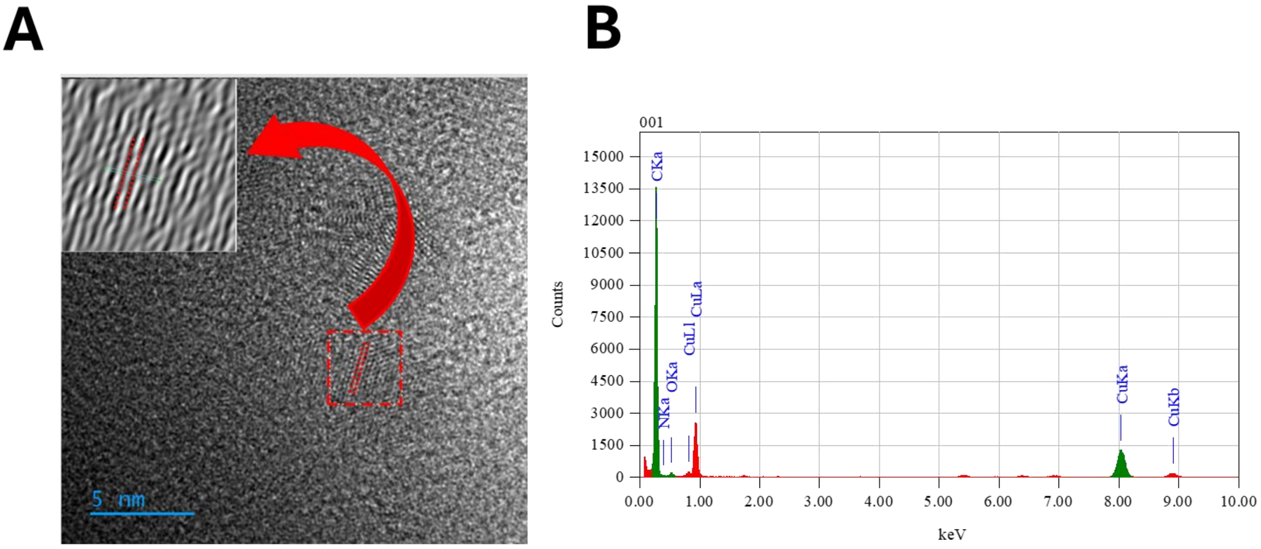


**Figure S3.** A. HRTEM image of Pap-CuNC. IFFT image of (inset) gives a lattice spacing of 0.2065±0.01 nm corresponding to the (111) plane of $Cu^0$. B. TEM-Energy Dispersive X-ray Spectra (EDS) of Pap-CuNC showing the presence of Copper

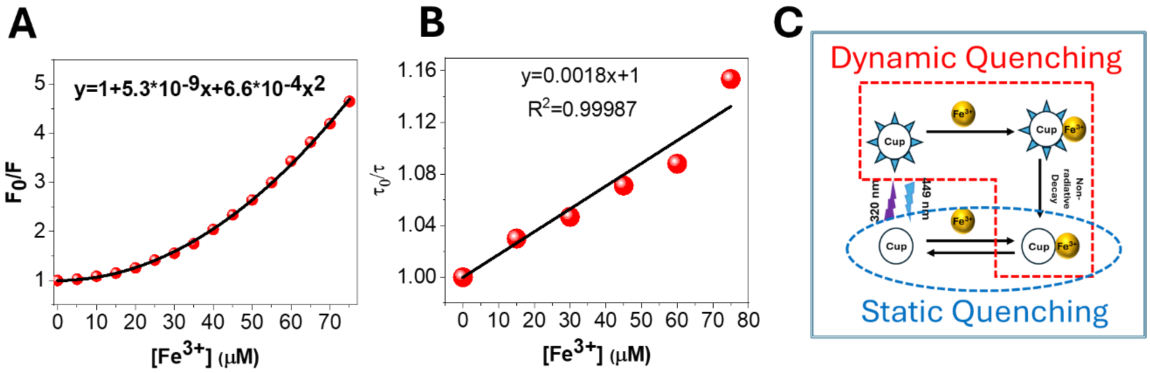


**Figure S4. Selective detection of Ferric ions by Pap-CuNC :** A. Stern-Volmer plot of intensity corresponds to the quenching of fluorescence intensity of Pap-CuNC by gradual addition of $Fe^{3+}$ ions. The black line represents second-order polynomial fitting of the parabolic pattern B. Stern-Volmer plot of lifetime corresponds to the quenching of fluorescence lifetime of Pap-CuNC by gradual addition of $Fe^{3+}$ ions. The black line represents linear fit C. A schematic mechanics for combine static and dynamic quenching

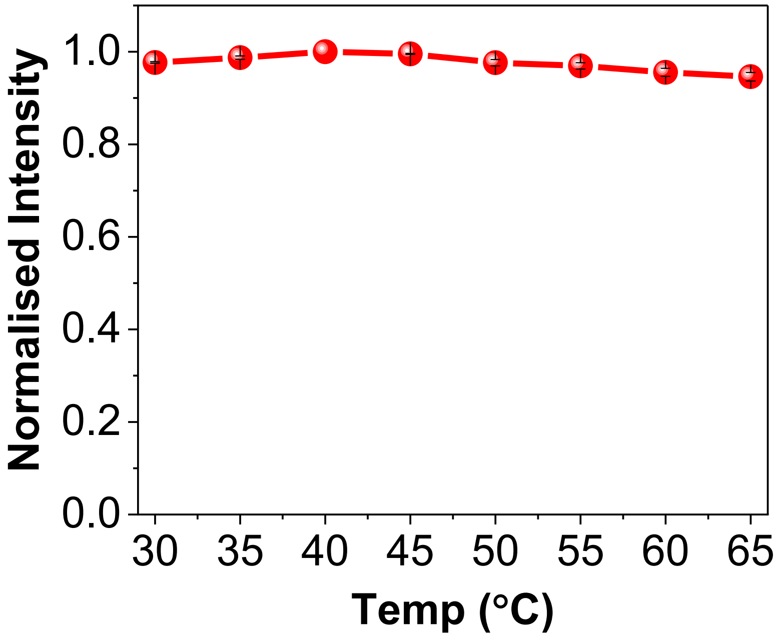


**Figure S5.** The emission intensity of Pap-CuNC on emission maxima does not change in a physiologically relevant temperature range (30-50°C)

**Video S1.** CuNC binding to the active site of the papain enzyme
